# Genomic Characterization of the RyC collection: 50 Multidrug Resistant Clinical Isolates of *Escherichia coli* and *Klebsiella* spp.

**DOI:** 10.64898/2026.05.18.725816

**Authors:** Paloma Rodera-Fernandez, Jorge Sastre-Dominguez, Coloma Costas, Javier DelaFuente, Aida Alonso-del Valle, Marta Hernández-García, Rafael Cantón, Alfonso Santos-Lopez, Alvaro San Millan

**Affiliations:** Centro Nacional de Biotecnología, Consejo Superior de Investigaciones Científicas (CSIC), Madrid, Spain; Departamento de Biología Molecular, Facultad de Biología, Universidad Autónoma de Madrid, Madrid, Spain; Instituto de Biología Evolutiva, Consejo Superior de Investigaciones Científicas (CSIC), Barcelona, Spain; Servicio de Microbiología. Hospital Universitario Ramón y Cajal and Instituto Ramón y Cajal de Investigación Sanitaria, Madrid, Spain; Centro de Investigación Biomédica en Red de Enfermedades Infecciosas (CIBERINFEC), Instituto de Salud Carlos III, Madrid, Spain; Centro de Investigación Biomédica en Red de Epidemiología y Salud Pública (CIBERESP), Madrid, Spain

**Keywords:** Enterobacterales, antimicrobial resistance, multidrug resistance, *Klebsiella pneumoniae*, *Escherichia coli*, plasmid, mobile genetic elements, pangenome, resistome, mobilome

## Abstract

Antimicrobial resistance (AMR) is a major global public health threat, and Enterobacterales producing extended-spectrum β-lactamases (ESBLs) represent some of the most common and concerning pathogens in clinical settings. Importantly, the dissemination of these resistance mechanisms is largely driven by mobile genetic elements (MGEs), particularly plasmids. Advancing our understanding of AMR evolution through experimentation requires moving beyond domesticated laboratory strains and towards clinically relevant isolates. However, despite the abundance of genomic data in public repositories, there is a lack of well-characterised clinical collections available for experimental work. Here, we characterise the RyC collection, which includes 50 multidrug-resistant, ESBL-producing *Escherichia coli* and *Klebsiella* spp. strains isolated from the gut microbiota of hospitalised patients at Hospital Universitario Ramón y Cajal (Madrid, Spain). We generated high-quality genome assemblies for all strains using a combination of short- and long-read sequencing technologies. From these data, we performed a comprehensive characterisation of the pangenome, mobilome, resistome and defensome of the collection. We present the RyC collection as a robust and experimentally tractable resource to study AMR evolution and MGEs dynamics in clinically relevant bacterial backgrounds.

**Impact statement:** Antimicrobial resistance (AMR) is a growing global health threat driven by the rapid dissemination of resistance genes among clinically relevant bacteria. A major challenge in studying AMR evolution is the reliance on domesticated laboratory strains, which poorly represent the complexity of pathogens circulating in hospitals. Here, we introduce the RyC collection, a set of well-characterised, multidrug-resistant Enterobacterales isolates obtained from hospitalised patients. By combining high-quality genome sequencing with detailed analyses of their gene content and mobile genetic elements (MGEs), this collection provides a realistic and experimentally tractable system to study how resistance evolves and spreads. The RyC collection will facilitate research on AMR dynamics, plasmid biology and host–MGEs interactions, ultimately contributing to the development of more effective strategies to combat antibiotic-resistant infections.

## Introduction

Antimicrobial resistance (AMR) has become a major global health concern, particularly in clinical settings where infections are becoming increasingly difficult to treat [1–3]. Extensive antibiotic use has accelerated the emergence of multidrug-resistant (MDR) isolates, limiting therapeutic options [4–6]. Enterobacterales stand out among the top antimicrobial resistant nosocomial pathogens prioritised by the WHO [7]. Specially worrisome are carbapenemase-producing Enterobacterales (CPE) and extended–spectrum ß-lactamase (ESBL)-producing Enterobacterales, which synthetise enzymes that degrade last-resort antibiotics widely used in MDR infections (e.g. carbapenems and third generation cephalosporins, respectively). Within CPE and ESBL-producing Enterobacterales, certain clones of *Klebsiella pneumoniae* and *Escherichia coli* represent the highest clinical threats, mainly due to their enhanced capacity to disseminate within hospital settings [8].

The pervasive dissemination of AMR, and consequently, the success of resistant pathogens in clinical environments is largely driven by mobile genetic elements (MGEs). MGEs enable the horizontal transfer of antimicrobial resistance genes (ARGs) across bacteria and play a pivotal role in AMR evolution [9, 10]. Among these elements, plasmids serve as critical vehicles of resistance, often carrying ARGs and virulence factors that can spread rapidly between strains. As a result, conjugative plasmids have been central to the emergence of globally successful MDR lineages [11–13]. Alongside plasmids, other MGEs such as transposable elements, phages, and integrons collectively contribute to the intricate mobilome (i.e. the collection of MGEs in a genome) that drives the adaptation of pathogens [14]. Understanding MGEs as interconnected systems is essential because their interactions often determine the stability and mobility of resistance and virulence factors [15]. Therefore, these elements not only spread resistance genes but also influence the general ecological and evolutionary success of bacterial pathogens [16, 17].

In this work, we present the complete genomic characterization of 50 relevant clinical strains of ESBL-producing Enterobacterales isolated during a large-scale surveillance program called R-GNOSIS [18]. The R-GNOSIS collection was assembled at the *Hospital Universitario Ramón y Cajal* (HURyC, Madrid, Spain) during a screening program for ESBL/carbapenemase-producing Enterobacterales colonising the gut microbiota of hospitalised patients. The program included the analysis of 28,089 samples obtained from 9,275 hospitalized patients between 2014 and 2016. Out of this collection, in previous research projects we selected 25 ESBL-producing *E. coli* and 25 ESBL-producing *Klebsiella spp.* representative of the phylogenetic diversity in the hospital. We then generated an isogenic collection by conjugating plasmid pOXA-48 into these strains (i.e. genetically identical except for the pOXA-48 presence). pOXA-48 is a worldwide spread PTU L/M carbapenem resistance plasmid highly prevalent in the HURyC [19, 20]. This collection of strains, which we name here the RyC collection, has been extremely informative in previous studies from our laboratory. For instance, we used all or part of the RyC collection to assess plasmid fitness effects [19–21], plasmid-chromosome transcriptomic crosstalks [22], or novel mechanisms of AMR evolution [23, 24] (**S. Table 1**). In this study, we performed long-read sequencing and hybrid genome assembly to comprehensively characterize the pangenome, mobilome and resistome of this collection. We expect that this thoroughly characterized collection of clinical strains will be of high value for the wider research community.

## Results

### The RyC collection

We previously selected 50 wild-type ESBL-producing Enterobacterales strains (25 *E. coli* and 25 *Klebsiella* spp.) that were representative of the diversity of the original R-GNOSIS collection [18, 19] (see methods for more details). Next, we conjugated pOXA-48 into these strains, generating isogenic pairs. All 100 clones underwent Illumina short-read sequencing for prior lab studies [19]. For the analyses presented in this work, we additionally sequenced the whole genomes of the 50 pOXA-48-free, wild type strains using Oxford Nanopore Technologies (ONT) long-read sequencing (See *Methods*). We then performed the hybrid assembly of the genomes using both short and long reads to obtain high quality closed genomes (**Fig. 1A**). Due to the complexity of clinical strains, achieving complete genome closed assemblies required manual curation of some strains and polishing to resolve ambiguities and ensure structural accuracy (see **Supp. Table 1**).

**Figure 1.**
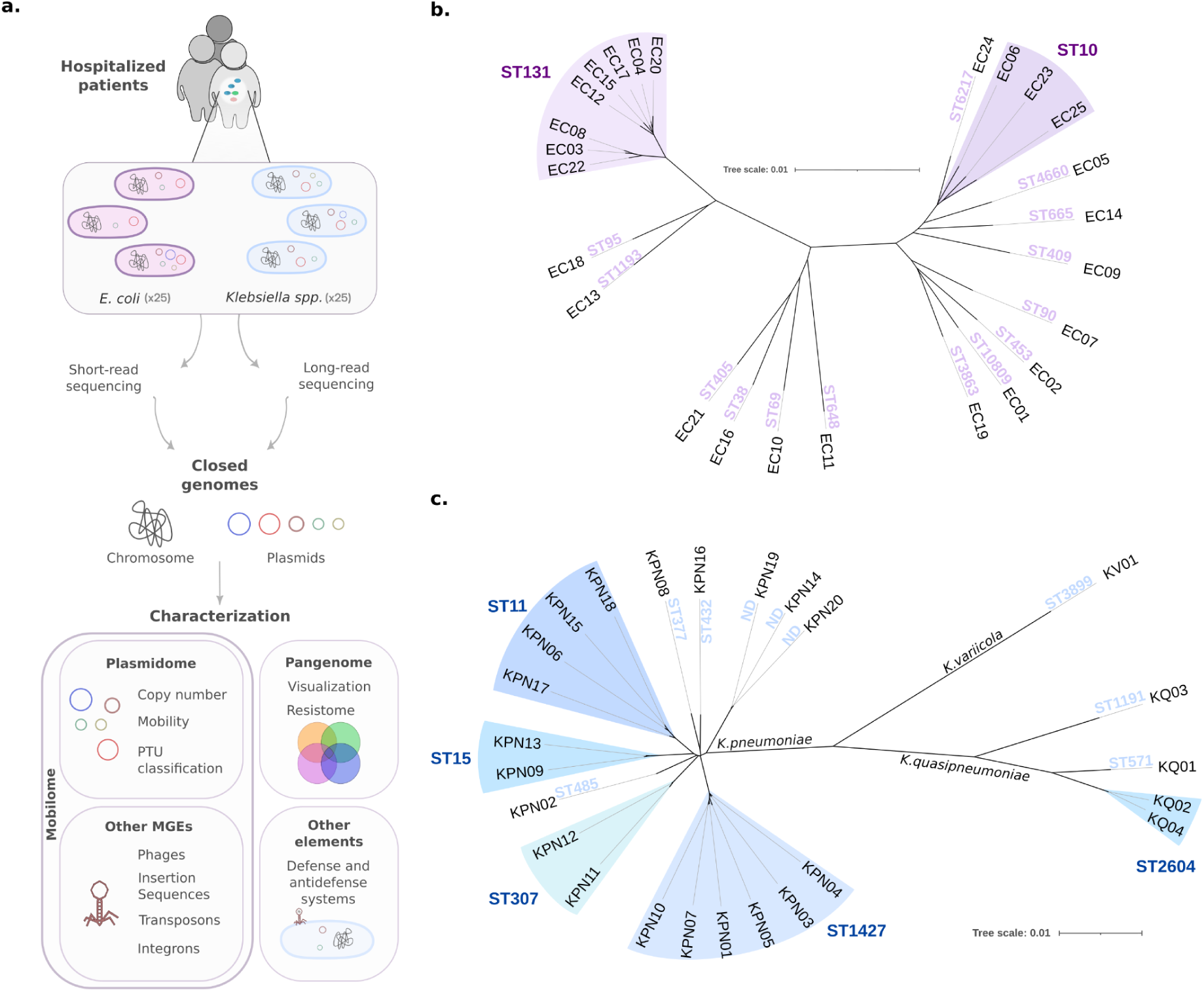
Characterization of the RyC collection. **A)** Schematic representation of the workflow used in this study. We selected 25 *E. coli* and 25 *Klebsiella* spp. isolated from the gut of hospitalized patients and sequenced them by short-read (Illumina) and long-read (Oxford Nanopore) technologies. We hybrid assembled the whole genome of each strain and characterized their pangenomes. Then, we analysed the strains’ mobilome which includes plasmids, phages, integrons and ISs. We also characterized other relevant elements for the bacterial mobilome such as defense and antidefense systems. **B)** Unrooted Neighbour Joining phylogenetic tree of the whole-genome assemblies from the 25 clones of *E. coli* characterized in this study. The grouping of multilocus sequence types (MLST) is highlighted, with the main ST groups being ST131 and ST10 (*E. coli* ST6217 belongs to the ST10 group). The scale represents mash distances between whole-genome assemblies. **C)** Unrooted Neighbour Joining phylogenetic tree of whole-genome assemblies from the 25 samples of *Klebsiella spp.* characterized in this study. The grouping of MLST is also highlighted, with the main ST groups being ST11, ST15, ST307, ST1427 and ST2604. The scale represents mash distances between whole-genome assemblies.

The RyC collection includes clones of ESBL-producing Enterobacterales of critical importance in clinical settings, belonging to four different species (*E. coli*, *n* = 25; *K. pneumoniae*, *n* = 20; *K. quasipneumoniae*, *n* = 4; *K. variicola*, n = 1, **Supp. Table 1**). For *E. coli*, eight isolates belonged to the sequence type (ST) ST131, a globally disseminated pathogenic lineage frequently associated with MDR infections, particularly prevalent in long-term care facilities [25, 26]. Three isolates belonged to ST10, another high-risk lineage prioritized by the WHO, characterized by their AMR and environmental persistence [27, 28]. Each of the remaining *E. coli* strains belonged to different STs, many of which were clinically relevant [29–31]. *K. pneumoniae* (**Fig. 1C**) strains belonged to different STs of clinical relevance, such as ST11 (n = 4), ST15 (n = 2), ST307 (n = 2) and ST1427 (n = 6). These STs are well-established high-risk carbapenem-resistant (CRKP) and MDR lineages with increasing prevalence in healthcare settings [32–36]. The remaining *Klebsiella* spp. strains belonged to individual STs (n = 6) or to unknown STs which were not previously described (annotated as ND, n = 3, KPN14, KPN19 and KPN20).

### Pangenome and resistome analysis

To assess the overall gene content within *E. coli* and *Klebsiella spp*. lineages, we conducted a pangenome analysis for each genus using Roary for construction and Anvi’o for visualisation [37, 38]. The resulting *E. coli* pangenome comprised 13,875 genes, of which 2,957 formed the core genome, present in all 25 isolates (**Fig. 2A**). An additional 199 genes represented the soft core (found in 95–99% of the genomes); the shell genome (found in 15–95% of the genomes) included 3,485 genes, while the remaining 7,234 genes—present in fewer than 4 strains—constituted the cloud genes (found in < 15% of the genomes). The *Klebsiella spp*. pangenome included 13,315 genes, with 3,182 shared across all isolates, defining the core genome (**Fig. 2B**). 261 genes formed the soft core genome and 3,447 genes the shell genome, while the remaining 6,425 were part of the cloud genome.

**Figure 2.**
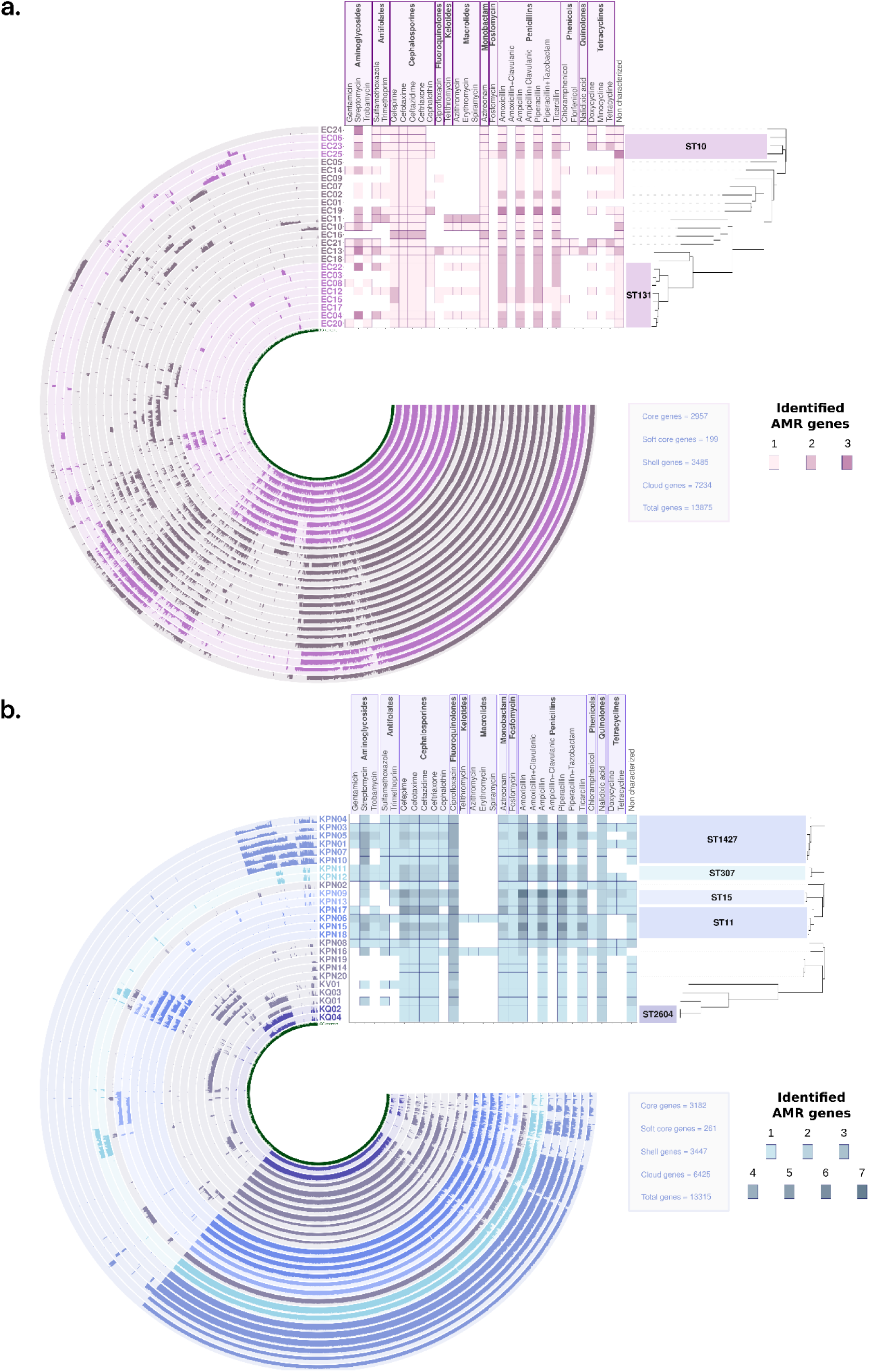
Pangenome and resistome of the RyC collection. Pangenome of the 25 *E. coli* (**A)** and 25 *Klebsiella spp.* (**B**) strains present in our collection, ordered by their phylogenetic distance tree. The main identified ST groups are represented in different tones of pink (*E. coli*) and blue (*Klebsiella spp.*). Information about the number of genes belonging to each category within the pangenome can be found in the grey box, on the right side of each pangenome. Heatmaps represent the AMR genes distribution (resistome) detected by Abricate using the Resfinder database [39]. The x-axis represents the different antibiotics to which the strains were predicted to be resistant and the y-axis, the strains.

Next, we characterized the ARGs, i.e. the resistome, of the collection using Resfinder (all the information about the ARGs, their location and the antibiotic to which they confer resistance can be found in **Supp. Table 2**) [39, 40]. Out of the 480 ARGs copies detected in the entire collection, 66% were encoded in plasmids (**Supp. Fig. 1**), while the rest were encoded in the chromosome. In only 5 cases the same ARG was found both in the chromosome and in plasmids in the same strain, *bla*_TEM-1D_ in one strain (EC19) and *bla*_CTX-M-15_ in 4 different strains (KPN05, KPN15, KPN17, KPN18). We identified 69 distinct AMR genes across our collection (**Supp. Fig. 2**), and we predicted the ARGs to confer resistance to 12 different classes of antibiotics (**Fig. 2AB**). Specifically *E. coli* presented resistance to 30 antibiotics and *Klebsiella spp*. to 28. Every strain was predicted to be resistant to at least 3 distinct classes of antibiotics (cephalosporins, monobactams and penicillins), and specifically to 9 antibiotics: amoxicillin, ampicillin, aztreonam, cefepime, cefotaxime, ceftazidime, ceftriaxone, piperacillin and ticarcillin, confirming that all of our strains present MDR genotypes [41]. The number of antibiotics to which each strain encoded putative resistance ranged from 9 in EC01, to 26 in KPN06. Overall, we detected predicted resistance to ß-lactams in 100% of the strains, to aminoglycosides in 74%, to antifolates in 64%, and to fluoroquinolones in 58%. To confirm the results obtained with Resfinder, we repeated the analysis using AMRFinderPlus (**Supp. Fig. 3** and **Supp. Fig. 4**), and obtained consistent results [42].

**Figure 3.**
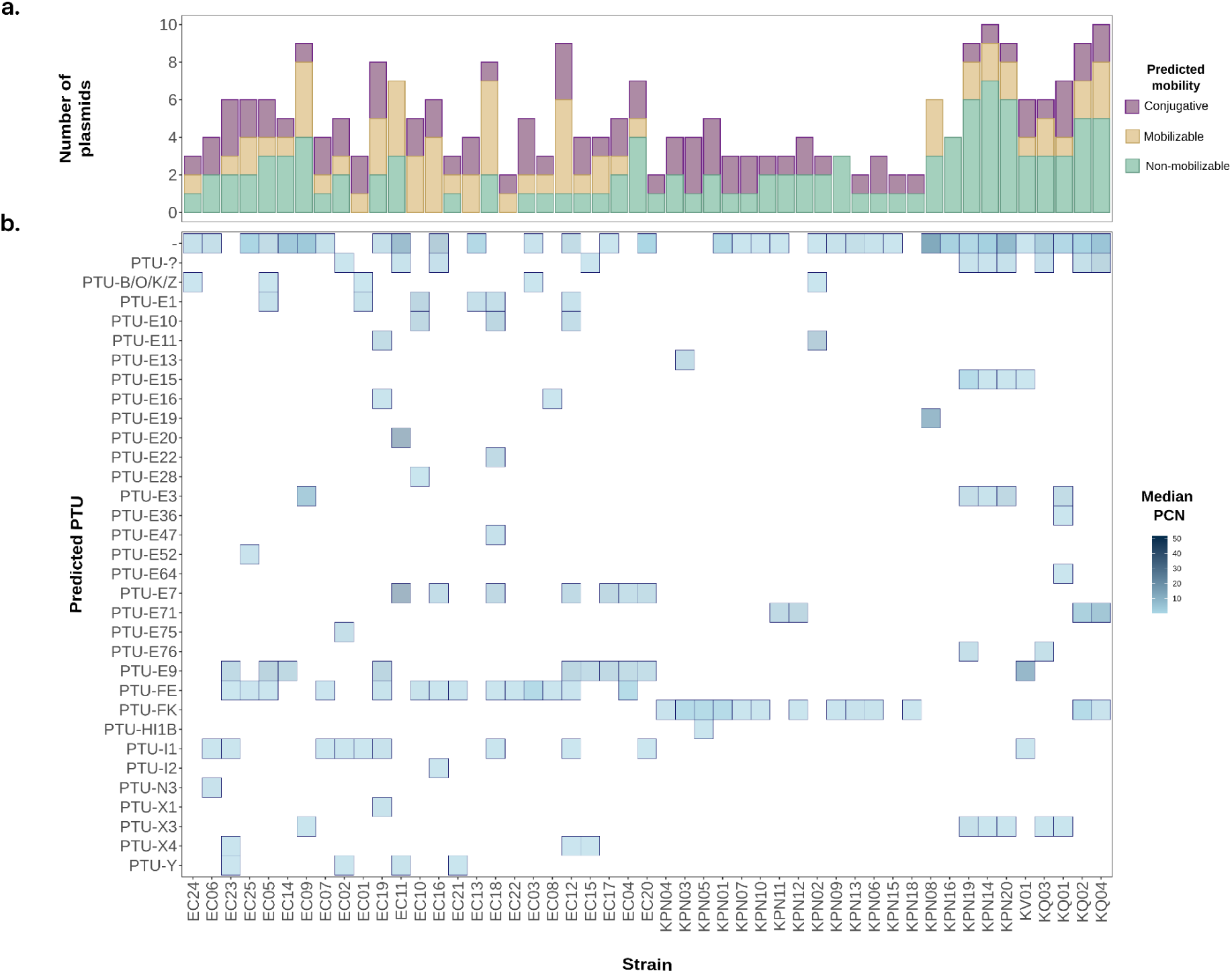
Characterization of the plasmidome of the RyC collection. a) Number of plasmids per strain. The x-axis represents all the strains of the collection (labelled in panel b), ordered according to the phylogenetic tree. The y-axis represents the number of plasmids per strain, differentiating by their predicted mobility. Conjugative plasmids are represented in purple, mobilizable plasmids in yellow and non-mobilizable plasmids in green. b) Heatmap showing the predicted PTUs per strain with its PCN. Note that new, unclassified PTUs are labeled as “PTU-?” and not identified PTUs are labeled as “-”, usually correlating to small and incomplete plasmids. The x-axis represents all the strains of the collection in the same order as in panel a. The y-axis represents the PTUs predicted by COPLA. The intensity of blue shade represents the PCN of each plasmid calculated as the plasmid’s median sequencing coverage divided by the chromosome’s median sequencing coverage.

**Figure 4.**
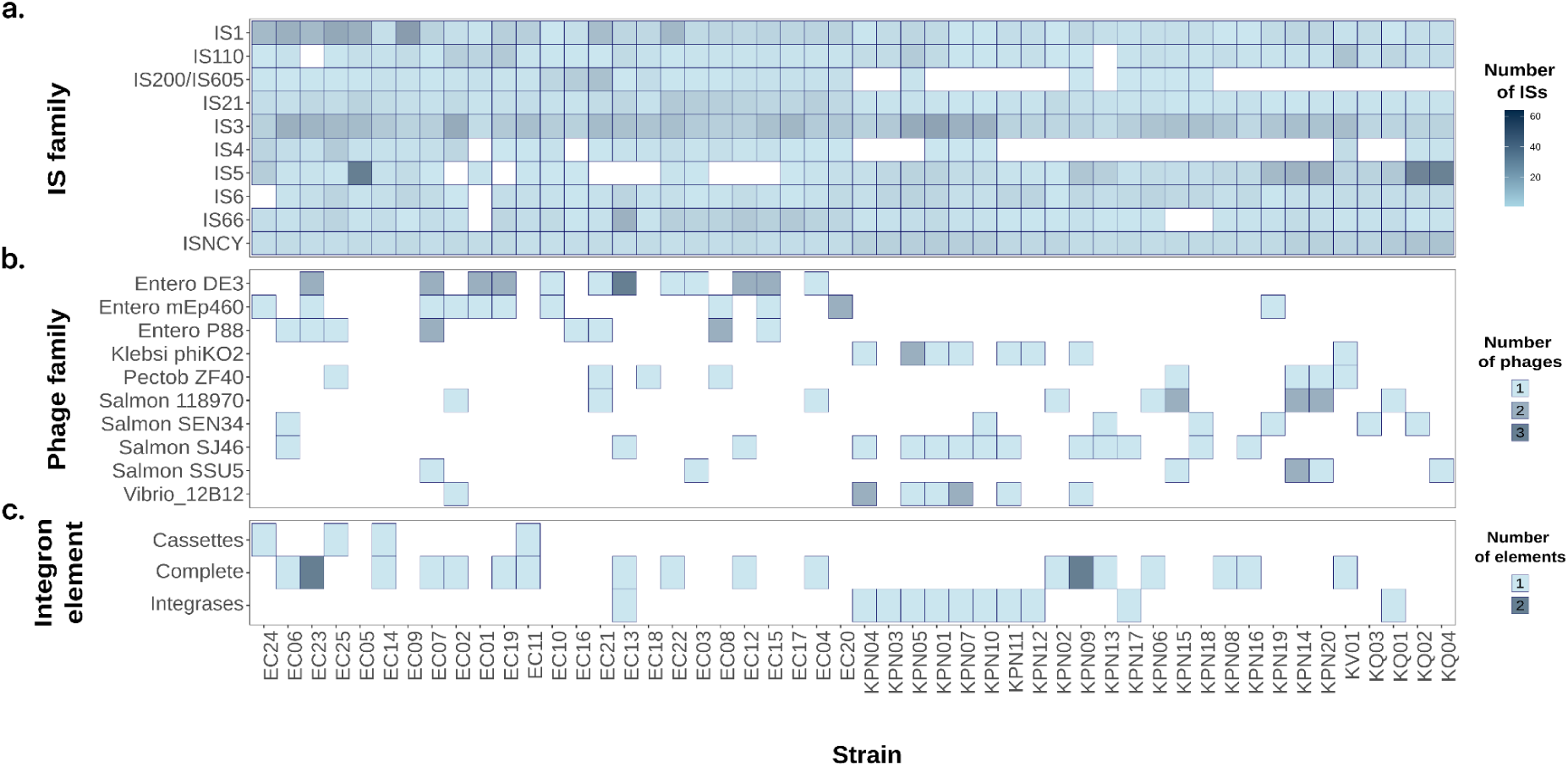
Distribution of ISs, prophages and integrons in the RyC collection. **I.** A) Heatmap of the IS families distribution across the collection. The y-axis displays the ten most prevalent IS families, with shade indicating the number of copies of each family per strain. The x-axis represents all the strains of the collection (labelled in panel C) ordered according to the phylogenetic tree. B) Heatmap showing the prophage families distribution. The y-axis displays the ten most prevalent phage families, with shade indicating the number of copies of each family per strain. The x-axis represents all the strains of the collection. C) Heatmap showing the integrons elements distribution across the collection. The y-axis shows the different elements that constitute an integron, with colours indicating the number of elements detected. These components may include: the cassettes -defined by the attC sites and associated genes conforming the cassette-, complete integrons (integrase plus cassettes), or integrases -representing the integrase lacking cassettes-. The x-axis represents all the strains of the collection in the same order as in panel a and b.

### Mobilome analysis

Beyond plasmids, other MGEs also promote the capture, accumulation, rearrangement and spread of ARGs. Therefore, we conducted a comprehensive characterisation of the content of MGEs in our collection, including plasmids, prophages, ISs and integrons. Extended information about the mobilome can be found in **Supp. Table 2** (plasmidome) and **Supp. Table 3** (phages, ISs and integrons).

### Plasmids

We identified a total of 203 plasmids in our collection using MOB-typer [42]. Among them, 38.9% were predicted to be conjugative, 27.6% mobilizable and 33.5% non-mobilizable (**Fig. 3A**). We used the PlasmidFinder database with Abricate and COPLA to identify each replicon, combining an approach based on specific traits with one based on homology [40, 43, 44]. Overall, PlasmidFinder identifies 238 replicons (**Supp. Fig.5**), whereas COPLA detected 203 PTUs (**Fig. 3B**). This difference may be explained by the fact that COPLA relies on sequence similarity, which makes PTU assignment to incomplete plasmids more difficult and particularly affects small plasmids like ColRNA. COPLA assigned 31 known PTUs and several new, unclassified PTUs (designated as “PTU-?”), including 136 plasmids (66.9%). The remaining plasmids could not be assigned to a PTU, but component and cluster predictions are available in **Supp. Table 2**. Among the 31 PTUs, nine appeared to be exclusive to *Klebsiella* genus and 16 to *E. coli*. The most frequent PTUs were PTU-FK with 17 instances in *Klebsiella spp*. and PTU-FE with 16 in *E.coli*. On the other hand, PlasmidFinder identified 39 distinct replicons, nine exclusive to *E. coli* and 20 to *Klebsiella spp.* The most frequent replicons detected by PlasmidFinder were IncFIC and IncFII in *Klebsiella spp*. and ColRNAl1 in *E. coli*, each appearing 19 times. Lastly, we analyzed the PCN for each replicon based on COPLA data. The PTU with the highest copy number was PTU-E20, found in EC11 (median PCN ∼ 23). In total, 8 PTUs -including the former- could be classified as high-copy number plasmids (HCP > 10 copies).

**Figure 5.**
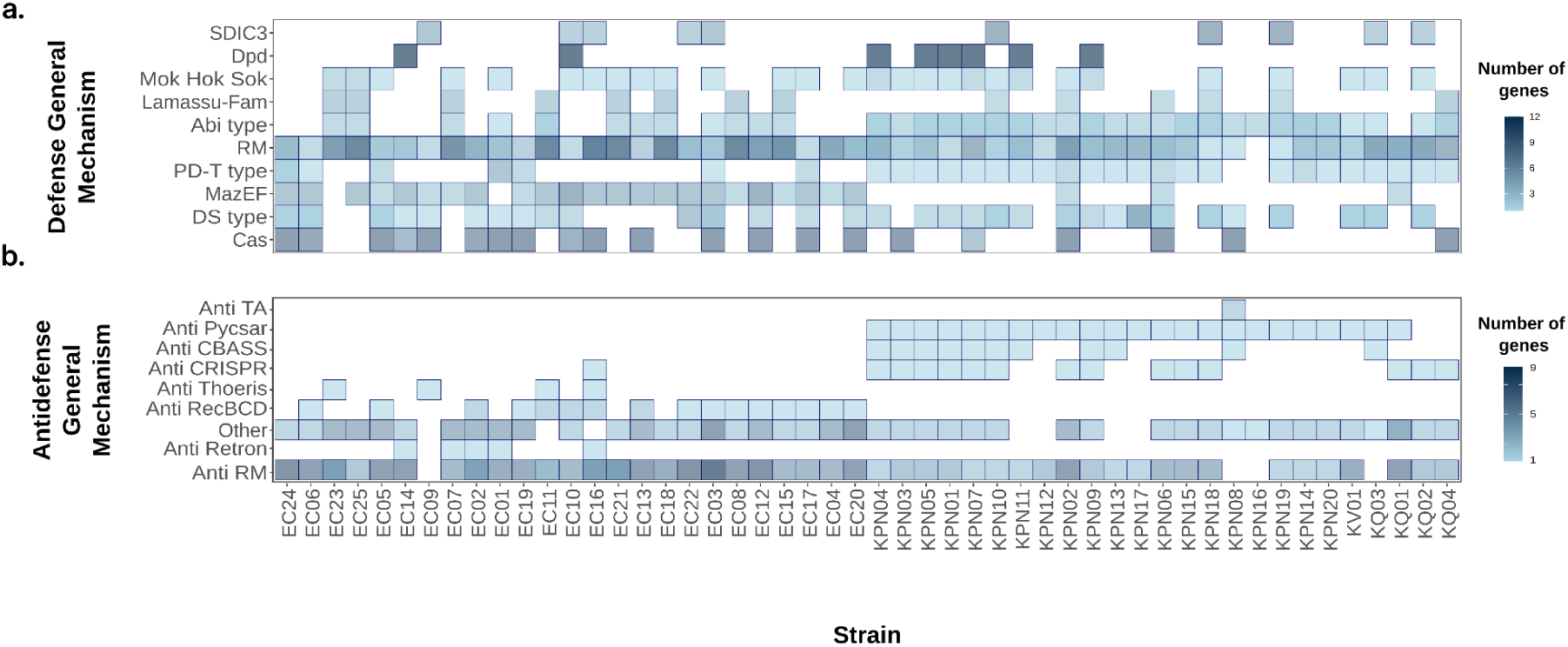
Defense/antidefense systems. A) Heatmap showing the defense systems across the collection. The y-axis displays the ten most prevalent defense mechanisms, with colour intensity indicating the number of copies of each family per strain. The x-axis represents all the strains of the collection (labelled in panel B) ordered according to the phylogenetic tree. B) Heatmap showing the antidefense systems across the collection. The y-axis displays the antidefense general mechanisms, with shade indicating the number of copies of each family per strain. The x-axis represents all the strains of the collection.

### IS elements

We detected a total of 4,285 IS copies across the collection belonging to 20 different IS families, 37.68% of them being encoded in plasmids (**Supp. Fig. 6**) [45]. The number of ISs per strain varied from 171 copies in EC05 to 41 in KPN02, with the IS5 being the most abundant element -up to 55 copies in EC05-, followed by the IS1 -up to 37 in EC09-. The IS1182 family appeared only in *E. coli* samples (EC11, EC14 and EC20, **Supp. Fig. 6**). The most prevalent IS families across the dataset were IS1, IS21 and IS3 families (**Fig. 4A**).

### Prophages

We identified prophages using the web-based-software PHASTEST, revealing a total of 302 prophages with varying levels of completeness (intact, questionable and incomplete) [46]. For downstream analysis, we retained only the 233 intact prophages, with a completeness score of above 90%, ensuring that we included only *bona fide* prophage sequences. To simplify the analysis, we retained from the original annotation only the main species and the precise name of the prophage, and selected the ten most prevalent prophage families (**Fig. 4B**). The number of distinct prophages ranged from 1 in KPN16, KQ02 and KQ04 to 10 in EC11, with 87% being encoded in the chromosome and 13% on plasmids (representing potential phage-plasmids) [47]. The phage with the highest copy number was the Enteric DE3, with 3 copies in EC16. This phage was *E. coli* specific and it’s the most common in our collection, appearing 20 times across 12 different samples [48, 49]. Another *E. coli* specific phage was the P88, described as P2-like coliphage [50].

### Integrons

We characterized integrons using Integron Finder, a tool that detects all the possible elements constituting an integron [51]. These include the complete integrons (integrase plus cassettes), the cassettes (defined by the *attC* sites and the genes contained in them), or only the integrase lacking associated cassettes. We found at least one integron-related element in 30 strains, with only 2 of them being encoded in the bacterial chromosome, and 28 encoded on plasmids (mobile integrons) (**Fig. 4C**). We identified complete integrons in 18 strains; most of them carrying just one, except for KPN09 and EC23, both carrying two complete integrons. The other identified elements were either the cassettes (n = 4) or the integrases (n = 9). We cross-referenced the cassettes positions with the Abricate results to detect cassettes encoding ARGs. We identified 1-3 ARGs across 24 integrons of varying completeness status. The most frequently identified genes were *aadA* (n = 17) and *ant(3”)-la* (n = 6), both associated with streptomycin resistance, and *dfrA* (n = 18), associated with trimethoprim resistance. Additionally, we identified genes associated with chloramphenicol (*cmlA;* n = 2) and sulfamethoxazole (*sul*1; n = 4) resistance.

### Defense and antidefense

Clinical Enterobacterales present a wide array of defense systems to control infection by MGEs. MGEs, in turn, encode a variety of antidefense systems to escape from bacterial immunity. To refine our analysis, we used Defense Finder to detect defense/antidefense systems in the strains [52–54]. All this information can be found in **Supp. Table 4**.

We first examined the defense mechanisms (**Fig. 5A**). We identified 617 defense systems across all the strains, 69.5% of them encoded in plasmids. Restriction-modification (RM) systems emerged as the most prevalent and abundant ones, with at least one RM present in all the strains except KPN17. Dpd, a recently reported anti-phage system, showed particularly high copy number, with 12 copies in two *E. coli* isolates and in 6 *K. pneumoniae*, being the system with the highest abundance per strain [55, 56]. Interestingly, we identified defense mechanisms originally associated with the *Vibrio* genus, such as VP1851 in KPN08 and Taranis in KQ01 (see **Supp. Table 4**). Recent studies have shown that some *Vibrio* defense systems are located specifically in the chromosome, near integrases, suggesting that they are part of MGEs [57, 58].

We next focused on the antidefense mechanisms identifying 345 systems, grouped into nine different types (**Fig. 5B**) [48, 59, 60]. Slightly more than a half of these systems (50.8%) were encoded in plasmids. The most widespread bacterial antidefense system is anti-RM, which prevents the formation of restriction barriers that block foreign DNA influx [61, 62]. Anti-RM genes were present in all strains except EC09, KPN08, KPN16 and KQ03, with up to nine copies found in EC03. Anti-CRISPR proteins were detected in 14 samples of *Klebsiella spp.*.

## Discussion

Despite the critical clinical importance of ESBL- and carbapenemase-producing Enterobacterales, the availability of high quality and comprehensively characterized collections of these strains for research purposes is scarce. In this study, we characterise the complete genomes of 50 multidrug resistant, clinically relevant strains of *E. coli* and *Klebsiella spp*. This collection includes clones of globally distributed high-risk clones, which were obtained from the gut microbiota of hospitalized patients in the *Hospital Universitario Ramón y Cajal* [18]. In addition to the 50 ESBL-producing Enterobacterales, this collection also includes a pOXA-48-carrying isogenic version of each strain. Therefore, this resource allows the in-depth analysis of both ESBL- and carbapenemase-producing strains. The RyC collection represents a useful experimental system for investigating AMR and MGEs dynamics. It provides a robust, high-quality framework to interrogate the genomic factors shaping the evolution and spread of AMR in clinical settings, as shown by previous works with these strains [19–24]. We expect that the RyC collection will represent a valuable experimental resource, and we make these strains accessible for the research community on demand.

To characterize the genomes of the RyC collection, we implemented a comprehensive bioinformatic workflow designed to describe AMR-associated genomic features in clinical Enterobacterales. Traditionally, genomic analyses of MDR Enterobacterales have often focused mostly on ARG-encoding plasmids, describing their distribution and mobility types. However, recent studies have increasingly highlighted the relevance of other MGEs, and their interactions with plasmids, to resistance evolution [63, 64]. Moreover, defense and antidefense systems also play a key role shaping MGEs dynamics and AMR evolution [52, 65]. Therefore, our workflow not only offers an in-depth plasmid analysis, but also integrates mobilome and defensome characterization using state-of-the-art bioinformatic tools. We hope that this workflow will be a useful asset for future genomic analyses of clinical Enterobacterales.

It is important to note that the workflow presented in this study relies on high quality genomic data obtained from hybrid assemblies which combine short (Illumina) and long (ONT) reads sequencing technologies. The use of this data is essential to comprehensively address the features related to MGEs in clinical bacteria (e.g. their genetic context, physiological features such as PCN, or the correct resolution of highly repetitive regions). One limitation of our workflow is that it relies on different softwares developed for the analysis of specific MGEs, some of which are only easily accessible through a web-based format. As a result, integrating and summarizing the results requires manual curation, which can be time-consuming. Therefore, there is a need for user-friendly software capable of analyzing the entire mobilome of bacterial genomes within an integrated framework. Such a tool would facilitate a deeper understanding of the genetic determinants underlying the success of clinically relevant bacteria.

## Methods

### Construction of the collection

The 50 strains used in this study were originally isolated as part of the R-GNOSIS collection and represent key clones within it. These strains were obtained through a surveillance screening program aimed at detecting extended-spectrum ß-lactamase (ESBL) and carbapenemase-producing Enterobacterales in hospitalized patients at the Ramón y Cajal University Hospital in Madrid, Spain, between 2014 and 2016 (Hernández-García et al., 2018; Maechler et al., 2020). The project, approved by the Hospital Ethics Committee (ref. no. 251/13), included a total of 28,089 samples collected from 9,275 patients hospitalized across four hospital wards: gastroenterology, neurosurgery, pneumology, and urology [18, 66]. The characterization of the samples was performed during the study period, rectal swabs were cultured on Chromo ID-ESBL and Chrom-CARB/OXA-48 selective agar media (BioMérieux, France). Colonies growing on these media were identified using MALDI-TOF MS (Bruker Daltonics, Germany) and further analysed by pulsed-field gel electrophoresis (PFGE). For the RyC collection, we selected 25 ESBL-producing *Escherichia coli* and 25 *Klebsiella spp.* isolates representing the diversity of these species, chosen randomly from the most prevalent PFGE profiles and ensuring the absence of the plasmid pOXA-48 [19]. The bacterial strains were cultured in LB broth at 37 °C with shaking at 250 rpm in 96-well plates, as well as on LB agar plates at 37 °C.

### Whole genome sequencing

#### Illumina short-read sequencing

We extracted the genomic DNA using the Wizard Genomic DNA Purification Kit (Promega Corporation). The short-read sequencing of these samples was performed at SeqCoast Genomics; at the Microbial Genome Sequencing Center (New Hampshire, USA) using the Illumina NextSeq200 platform. Paired-end short reads were 150 base pairs (bp) in length, with a mean coverage of 245.70.

#### Oxford Nanopore Technologies long-read sequencing

To generate closed assemblies for a proper genomic characterization, we sequenced the whole genome of the strains using Oxford Nanopore Technologies (ONT). We extracted the genomic DNA as explained in the section above. We then prepared the genomic libraries following the Nanopore protocol for native barcoding genomic DNA (EXP-NBD114.24 kit). For the quantification of the extracted DNA, we used a Qubit Flex Fluorometer (Thermo Fisher Scientific). To perform the DNA repair and end-prep steps, we used the NEBNext Ligation Sequencing Kit (Oxford Nanopore Technologies), AMPure XP beads (Beckman Coulter/Agencourt AMPure XP) and a HulaMixer (Thermo Fisher Scientific). For the native barcode ligation step, we used the EXP-NBD114.24 kit (Oxford Nanopore Technologies) and the Blunt/TA Ligase Master Mix (New England Biolabs). Then, we performed the adaptor ligation and clean-up steps with the NEBNext Quick Ligation Reaction Buffer (New England Biolabs), NEBNext Ligation Sequencing Kit (New England Biolabs). We carried out the priming and loading of the flow cell steps, in which we used the Flow Cell Priming Kit (EXP-FLP004) (Oxford Nanopore Technologies) and sequenced the samples using R10.4.1 Flow Cells (FLO-MIN114) in a MinION Mk1B (mean 23.20 coverage).

### Genome characterization of the collection

#### Quality control of Illumina reads

We carried out the quality control of the raw Illumina reads using FastQC v.0.12.1 and merged the reports using MultiQC v.1.27.1. We trimmed the reads using Trim Galore v.0.6.10, adapting the parameters to trim low-quality ends, discard reads shorter than 50 bp and to trim nextera adaptors (-q 20 –length 50 –nextera). We performed a second quality control of the trimmed reads and used them for the following downstream analyses.

#### *De novo* assembly of the genomes

We assembled the genomes of the strains for a proper identification of all the genetic features of the collection. We performed the hybrid assembly using both long (ONT) and short (Illumina) reads using Unicycler v.0.5.1 or Flye v.2.9.5-b1801 [68, 69]. We checked the assembly completeness using Bandage v.0.9.0 [70]. For those strains for which complete assembly could not be achieved, we performed manual curation of the partial assemblies using Bandage. All the detailed information of how we performed the assembly of each strain can be found in **Supp. Table 1**. We then annotated the assembled genomes with Prokka v.1.14.6 and determined the ST of all the isolates using the multilocus sequence-typing tool mlst v.2.23.0 [71, 72]. We generated the phylogenetic trees using the assemblies as an input for Mashtree 1.4.6 [73]. We visualized the phylogenetic trees using the iTOL platform v.7, integrating the ST information [74].

#### Pangenome generation

We generated the pangenome for each species using Roary v.3.13.0 using default parameters and the Prokka annotation [38]. We visualized the *E. coli* and the *Klebsiella spp.* pangenomes using Anvi’o v.8 by executing the microbial population workflow specified by the developers (https://merenlab.org/2016/11/08/pangenomics-v2/) [37]. Briefly, we mapped the clean reads against the pangenome using BWA-MEM v.0.7.19 and transformed the output into sorted and indexed BAM files using Samtools v.1.19.2 [75, 76]. We reformatted the assemblies using a specific program from Anvi’o to prevent incompatibilities. The annotation of the reformatted FASTA file was then repeated using Prokka, incorporating the previous annotation from PGAP (--proteins flag). Then, we extracted the gene and functional annotation using a script created by the Anvi’o developers. Afterwards, we created a reference database and imported the gene functions. To enhance visualization, we adjusted the parameters to enable the soft-splitting of large contigs, forcing the generation of splits with a desired length (length of 1,000bp), and to ignore amino acid sequences with stop codons. Lastly, we created a profile database for each sample, merged into a single profile and displayed the results.

#### Plasmidome and resistome characterization

To characterize the plasmids of all the strains in our collection, we applied multiple analytical methods. First, we ran Abricate v.1.0.1 using the Plasmidfinder (v.2024-Nov-19) and the ResFinder (v.2024-Nov-19) database to identify antimicrobial resistant determinants, both in the plasmid and the chromosomal genes [39, 40, 43]. To confirm the resistance predictions, we also used AMRFinderPlus, a software designed to detect ARGs, resistance-associated mutations, and virulence genes [77]. We selected this tool because it forms the basis of the ISO-certified workflow abritAMR [78]. We then compared the results obtained with ResFinder to those generated by AMRFinderPlus. Overall, 51.1% of the AMR genes were predicted in common by both databases. 19 genes were detected exclusively by ResFinder, including four different *bla*_SHV_ genes, three *bla*_TEM_ genes, and *mdf(A)*. In contrast, AMRFinderPlus identified 30 exclusive genes, including qacEdelta1. However, some of these discrepancies are likely due to differences in gene annotation. For example, oqxA in EC13 was annotated as *oqxA2* in AMRFinderPlus and as *oqxA_1* in ResFinder, and similar inconsistencies were observed for some *bla*_TEM_ genes. Second, we classified plasmids into incompatibility groups according to their replication mechanism, and predicted their mobility using MOB-suite v.3.1.9 with the –*multi* flag, which allows for the identification of independent plasmids within samples [42]. Finally, to overcome the limited precision of plasmid classification based solely on specific traits, we also ran COPLA v.1 [44]. This software classifies plasmids based on overall genetic similarity, grouping them into the same plasmid taxonomic unit (PTU) when they share a high homology across more than 50% of their sequence.

#### Identification of other mobile genetic elements and defense/antidefense systems

In order to thoroughly describe the genomes in our collection, we also performed an extensive analysis of relevant genomic determinants (defense and antidefense systems) and multiple mobile genetic elements (besides plasmids, including phages, IS and integrons). We used the web-based tool PHASTEST to identify viruses and proviruses across our sequences [46]. Then, we filtered and selected only the intact prophages, discarding those with score below 90. We used ISescan v.1.7.2.3 to detect IS elements, annotating their completeness and the number of copies present in each assembly [45]. We used Defense-finder v.2.0.1 to identify known anti-phage and anti-defense systems [53, 54, 79]. We used Integron-Finder with default parameters to identify the integrons and their composition.

## Author Notes

All supporting data, code and protocols have been provided within the article or through supplementary data files. Five supplementary figures and four supplementary tables are available with the online version of this article.

## Abbreviations

AMR: antimicrobial Resistance
ARG: antimicrobial resistance genes
CPE: carbapenemase-producing enterobacterales
CRKP: carbapenem-resistant *Klebsiella pneumoniae*
EC: *Escherichia coli*
ESBL: extended–spectrum ß-lactamase
HCP: high copy number
HURyC: *Hospital Universitario Ramón y Cajal*
RyC: Ramon y Cajal
IS: insertion sequence
KPN: *Klebsiella pneumoniae*
KQ: *Klebsiella quasipneumoniae*
KV: *Klebsiella variicola*
MDR: multidrug resistant
MGE: mobile genetic element
MLST: multilocus sequence type
PCN: plasmid copy number
PTU: plasmid taxonomic unit
RM: restriction-modification
ST: sequence type

## Data summary

All raw sequencing data used for this study are publicly available: PRJNA1455872. All custom code used in this article can be accessed at (https://github.com/Plasmidloma/RyC_collection).

## Code availability

The code developed for this project can be found in https://github.com/Plasmidloma/RyC_collection.

## Data availability

All the sequences generated for this project can be found at the Sequence Read Archive (SRA) repository of the National Center for Biotechnology Information (NCBI) under the BioProject ID PRJNA1455872.

## Contributions

A.S.M. acquired the funding and supervised the study. M.H.G and R.C contribute with the RYC collection and preliminary characterization through the R-GNOSIS project. A.A.-d.V., with help from J.D.F. and C.C., performed the experimental work. P.R.F. and J.S.D performed the bioinformatical and phylogenetic analysis. A.S.M., A.S.L, J.S.D and P.R.F. wrote the initial draft of the manuscript, and all the authors contributed to the final version of the manuscript and approved it.

## Supplementary figures

**Supplementary Figure 1.**
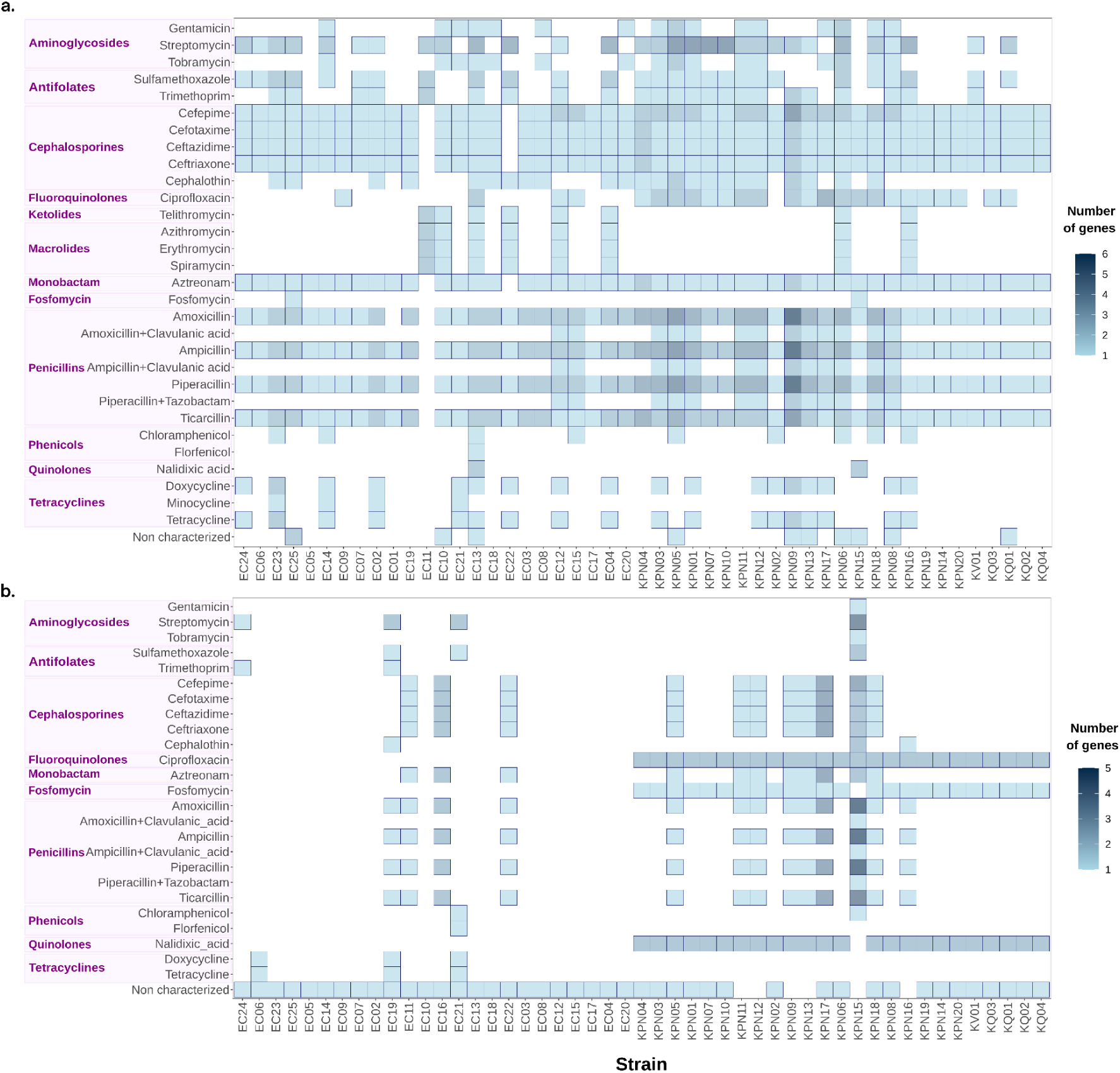
Heatmap showing the predicted antibiotic resistance encoded in plasmids (a) and in chromosome (b). The y-axis displays all the predicted antibiotics and their respective classes, with shade indicating the number of genes putatively conferring resistance to each class per strain. The x-axis represents all the strains of the collection ordered according to the phylogenetic tree.

**Supplementary Figure 2.**
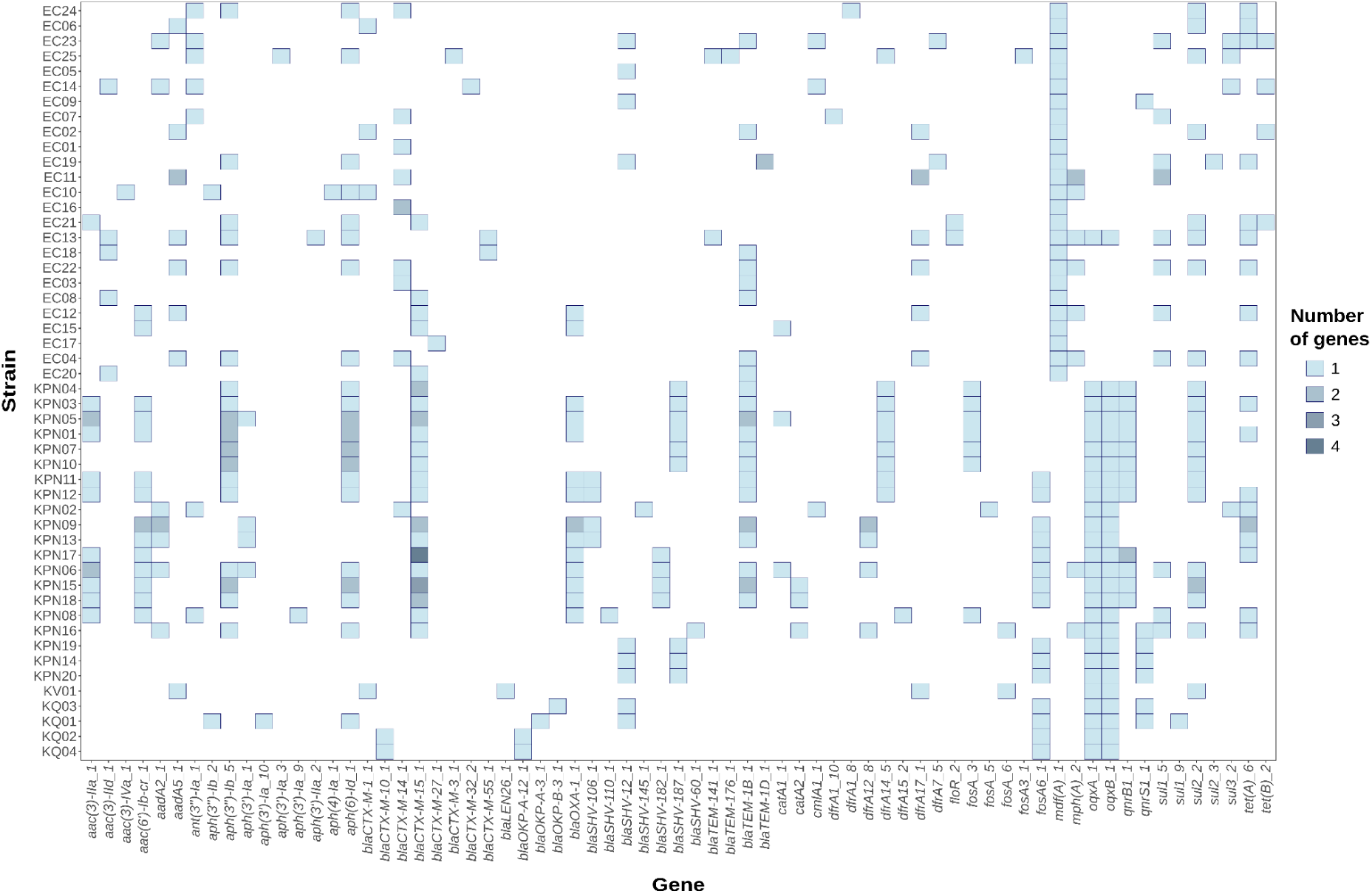
Heatmap showing the AMR genes detected by Resfinder. The y-axis represents all the strains of the collection ordered according to the phylogenetic tree. The x-axis displays all the AMR genes, with shade indicating the number of gene copies of each one per strain.

**Supplementary Figure 3.**
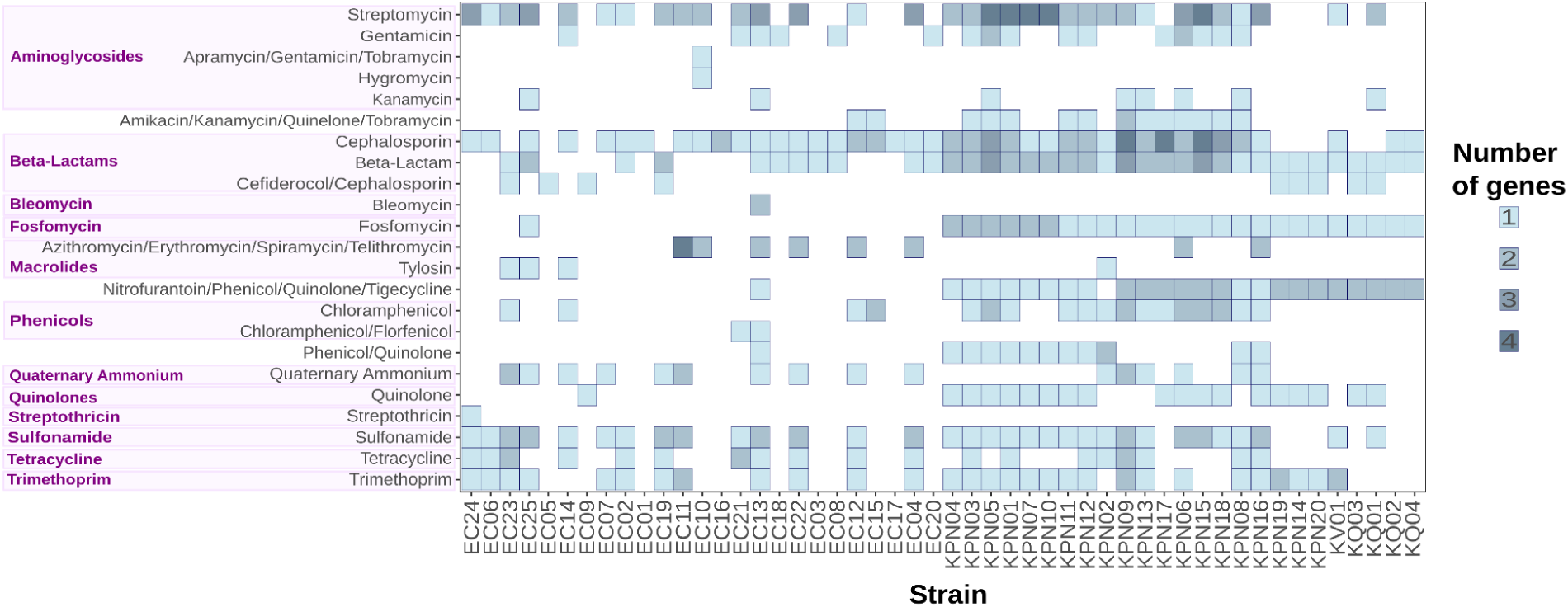
Predicted antibiotic resistance by AMRFinderPlus. Heatmap showing the predicted antibiotic resistance by AMRFinderPlus, a software for the identification of resistance genotype predictions. The y-axis displays all the predicted antibiotics and their respective classes, with shade indicating the number of genes putatively conferring resistance to each class per strain. The x-axis represents all the strains of the collection ordered according to the phylogenetic tree.

**Supplementary Figure 4.**
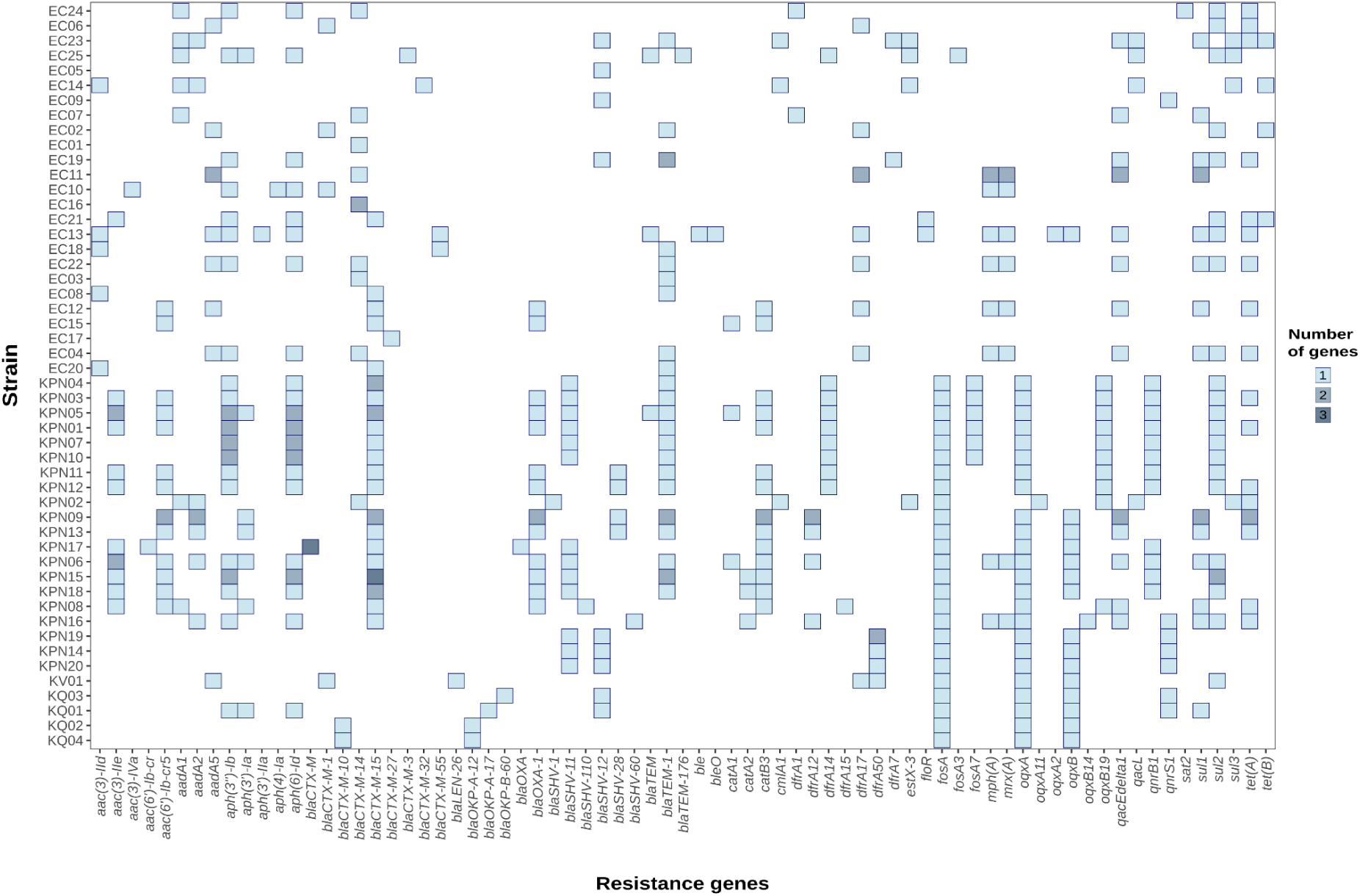
Heatmap showing the AMR genes detected by AMRFinderPlus. The y-axis represents all the strains of the collection ordered according to the phylogenetic tree. The x-axis displays all the AMR genes, with shade indicating the number of gene copies of each one per strain.

**Supplementary Figure 5.**
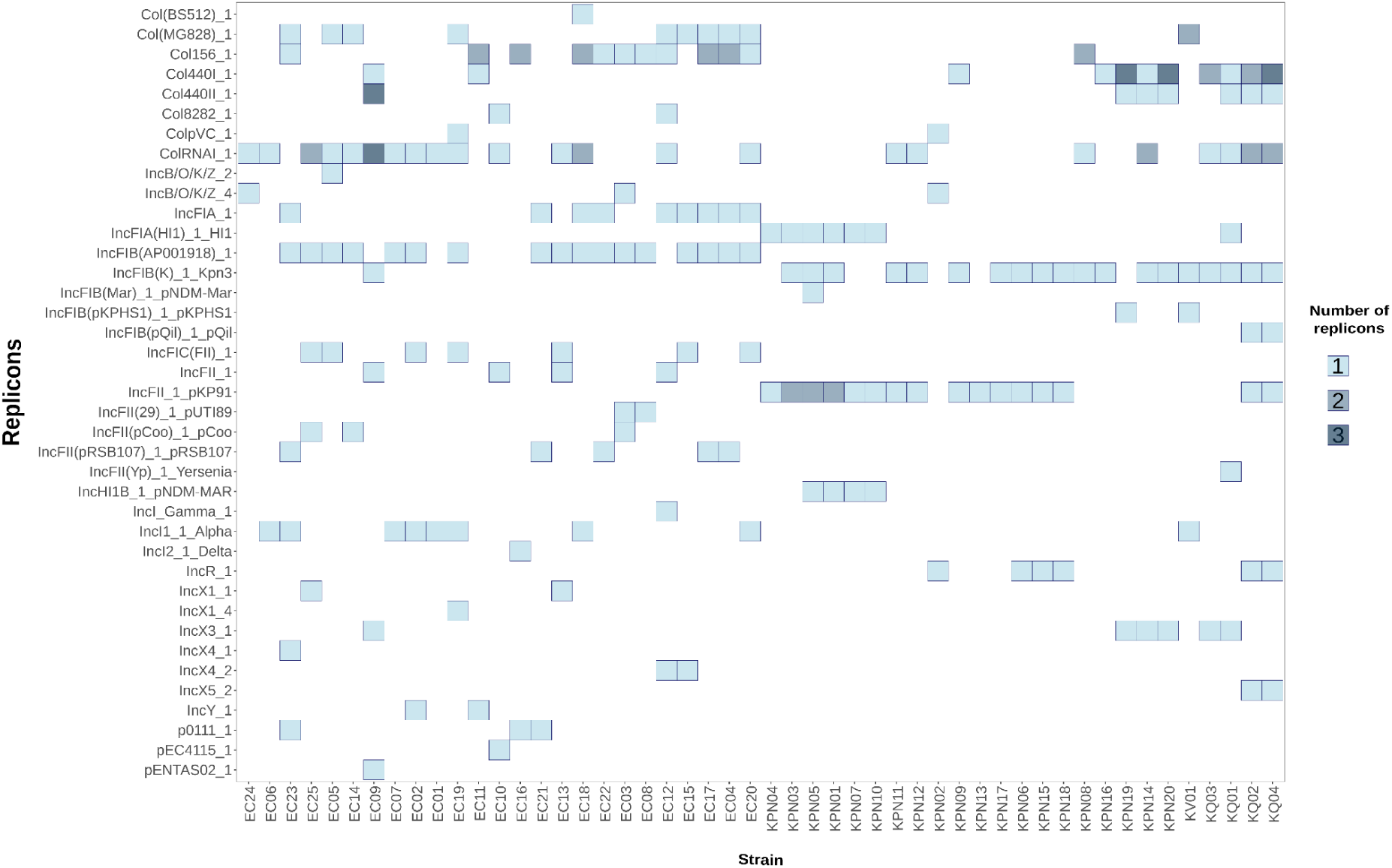
Replicon identification by Abricate. Heatmap of the replicons identified using PlasmidFinder across the collection. The y-axis displays all the replicons, with shade indicating the number of copies of each one per strain. The x-axis represents all the strains of the collection ordered according to the phylogenetic tree.

**Supplementary Figure 6.**
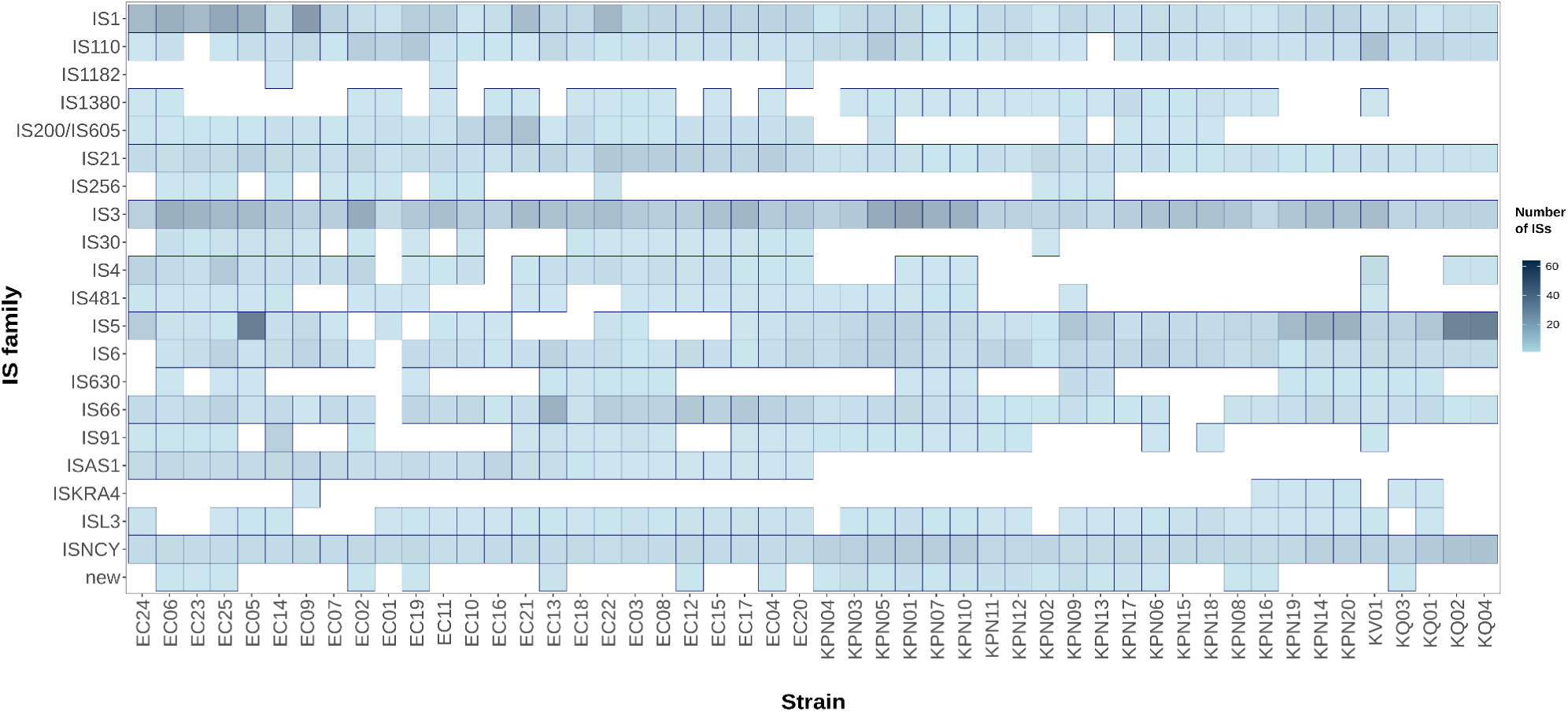
Distribution of ISs. Heatmap of the IS families distribution across the collection. The y-axis displays all the IS families, with shade indicating the number of copies of each family per strain. The x-axis represents all the strains of the collection ordered according to the phylogenetic tree.

## References

1. Ho CS, Wong CTH, Aung TT, Lakshminarayanan R, Mehta JS, et al. Antimicrobial resistance: a concise update. Lancet Microbe 2025;6:100947.

2. Kariuki S. Global burden of antimicrobial resistance and forecasts to 2050. The Lancet 2024;404:1172–1173.

3. Okeke IN, Kraker MEA de, Boeckel TPV, Kumar CK, Schmitt H, et al. The scope of the antimicrobial resistance challenge. The Lancet 2024;403:2426–2438.

4. Bassetti M, Righi E, Carnelutti A, Graziano E, Russo A. Multidrug-resistant Klebsiella pneumoniae: challenges for treatment, prevention and infection control. Expert Rev Anti Infect Ther 2018;16:749–761.

5. Kocsis B, Gulyás D, Szabó D. Emergence and Dissemination of Extraintestinal Pathogenic High-Risk International Clones of Escherichia coli. Life 2022;12:2077.

6. Lam MMC, Wick RR, Watts SC, Cerdeira LT, Wyres KL, et al. A genomic surveillance framework and genotyping tool for Klebsiella pneumoniae and its related species complex. Nat Commun 2021;12:4188.

7. Mukhopadhyay S, Peng Y, Tun HM. The 2024 WHO bacterial priority pathogens list: a critical evolution from a global One Health perspective. Sci One Health 2025;5:100145.

8. Bonomo RA, Burd EM, Conly J, Limbago BM, Poirel L, et al. Carbapenemase-Producing Organisms: A Global Scourge. Clin Infect Dis 2018;66:1290–1297.

9. Gogarten JP, Townsend JP. Horizontal gene transfer, genome innovation and evolution. Nat Rev Microbiol 2005;3:679–687.

10. MacLean RC, San Millan A. The evolution of antibiotic resistance. Science 2019;365:1082–1083.

11. Andrade LN, Curiao T, Ferreira JC, Longo JM, Clímaco EC, et al. Dissemination of blaKPC-2 by the Spread of Klebsiella pneumoniae Clonal Complex 258 Clones (ST258, ST11, ST437) and Plasmids (IncFII, IncN, IncL/M) among Enterobacteriaceae Species in Brazil. Antimicrob Agents Chemother 2011;55:3579–3583.

12. Naseer U, Sundsfjord A. The CTX-M Conundrum: Dissemination of Plasmids and Escherichia coli Clones. Microb Drug Resist 2011;17:83–97.

13. Valverde A, Cantón R, Garcillán-Barcia MP, Novais Â, Galán JC, et al. Spread of blaCTX-M-14 Is Driven Mainly by IncK Plasmids Disseminated among Escherichia coli Phylogroups A, B1, and D in Spain. Antimicrob Agents Chemother 2009;53:5204–5212.

14. Lang AS, Buchan A, Burrus V. Interactions and evolutionary relationships among bacterial mobile genetic elements. Nat Rev Microbiol 2025;23:423–438.

15. Pfeifer E, Bonnin RA, Rocha EPC. Phage-Plasmids Spread Antibiotic Resistance Genes through Infection and Lysogenic Conversion. mBio 2022;13:e01851–22.

16. Jackson RW, Vinatzer B, Arnold DL, Dorus S, Murillo J. The influence of the accessory genome on bacterial pathogen evolution. Mob Genet Elem 2011;1:55–65.

17. Richardson EJ, Bacigalupe R, Harrison EM, Weinert LA, Lycett S, et al. Gene exchange drives the ecological success of a multi-host bacterial pathogen. Nat Ecol Evol 2018;2:1468–1478.

18. Hernández-García M, Pérez-Viso B, Carmen Turrientes M, Díaz-Agero C, López-Fresneña N, et al. Characterization of carbapenemase-producing Enterobacteriaceae from colonized patients in a university hospital in Madrid, Spain, during the R-GNOSIS project depicts increased clonal diversity over time with maintenance of high-risk clones. J Antimicrob Chemother 2018;73:3039–3043.

19. Alonso-del Valle A, León-Sampedro R, Rodríguez-Beltrán J, DelaFuente J, Hernández-García M, et al. Variability of plasmid fitness effects contributes to plasmid persistence in bacterial communities. Nat Commun 2021;12:2653.

20. Alonso-del Valle A, Toribio-Celestino L, Quirant A, Pi CT, DelaFuente J, et al. Antimicrobial resistance level and conjugation permissiveness shape plasmid distribution in clinical enterobacteria. Proc Natl Acad Sci 2023;120:e2314135120.

21. Calvo-Villamañán A, Sastre-Dominguez J, Barrera-Martín Á, Costas C, San Millan Á. Dissecting pOXA-48 fitness effects in clinical Enterobacterales using plasmid-wide CRISPRi screens. Nat Commun 2025;16:7700.

22. Toribio-Celestino L, Calvo-Villamañán A, Herencias C, Alonso-del Valle A, Sastre-Dominguez J, et al. A plasmid-chromosome crosstalk in multidrug resistant enterobacteria. Nat Commun 2024;15:10859.

23. Sastre-Dominguez J, DelaFuente J, Toribio-Celestino L, Herencias C, Herrador-Gómez P, et al. Plasmid-encoded insertion sequences promote rapid adaptation in clinical enterobacteria. Nat Ecol Evol 2024;8:2097–2112.

24. Sastre-Dominguez J, Rodera-Fernandez P, DelaFuente J, Martínez-González S, Quesada S, et al. Plasmids promote antimicrobial resistance through insertion sequence-mediated gene inactivation. Nat Microbiol 2026;11:976–992.

25. Ludden C, Decano AG, Jamrozy D, Pickard D, Morris D, et al. Genomic surveillance of Escherichia coli ST131 identifies local expansion and serial replacement of subclones. Microb Genomics 2020;6:e000352.

26. Pitout JDD, Finn TJ. The evolutionary puzzle of *Escherichia coli* ST131. Infect Genet Evol 2020;81:104265.

27. Aibinu I, Odugbemi T, Koenig W, Ghebremedhin B. Sequence Type ST131 and ST10 Complex (ST617) predominant among CTX-M-15-producing *Escherichia coli* isolates from Nigeria*. Clin Microbiol Infect 2012;18:E49–E51.

28. Fuga B, Sellera FP, Cerdeira L, Esposito F, Cardoso B, et al. WHO Critical Priority Escherichia coli as One Health Challenge for a Post-Pandemic Scenario: Genomic Surveillance and Analysis of Current Trends in Brazil. Microbiol Spectr 2022;10:e01256–21.

29. Chattaway MA, Jenkins C, Ciesielczuk H, Day M, DoNascimento V, et al. Evidence of Evolving Extraintestinal Enteroaggregative Escherichia coli ST38 Clone. Emerg Infect Dis 2014;20:1935–1937.

30. Pitout JDD. Extraintestinal Pathogenic Escherichia coli: A Combination of Virulence with Antibiotic Resistance. Front Microbiol 2012;3:9.

31. Pitout JDD, Peirano G, Chen L, DeVinney R, Matsumura Y. Escherichia coli ST1193: Following in the Footsteps of E. coli ST131. Antimicrob Agents Chemother 2022;66:e00511–22.

32. Lam MMC, Wyres KL, Wick RR, Judd LM, Fostervold A, et al. Convergence of virulence and MDR in a single plasmid vector in MDR Klebsiella pneumoniae ST15. J Antimicrob Chemother 2019;74:1218–1222.

33. Peirano G, Chen L, Kreiswirth BN, Pitout JDD. Emerging Antimicrobial-Resistant High-Risk Klebsiella pneumoniae Clones ST307 and ST147. Antimicrob Agents Chemother 2020;64:10.1128/aac.01148-20.

34. Yang Y, Qin J, Hu Y, Wang J, Feng Y, et al. Carbapenem-Resistant Klebsiella pneumoniae of Sequence Type 11: A Scoping Review. J Infect Dis 2026;233:S72–S80.

35. Zhao J, Liu C, Liu Y, Zhang Y, Xiong Z, et al. Genomic characteristics of clinically important ST11 *Klebsiella pneumoniae* strains worldwide. J Glob Antimicrob Resist 2020;22:519–526.

36. Zhou K, Lokate M, Deurenberg RH, Arends J, Lo-Ten Foe J, et al. Characterization of a CTX-M-15 Producing Klebsiella Pneumoniae Outbreak Strain Assigned to a Novel Sequence Type (1427). Front Microbiol;6. Epub ahead of print 10 November 2015. DOI: 10.3389/fmicb.2015.01250.

37. Eren AM, Esen ÖC, Quince C, Vineis JH, Morrison HG, et al. Anvi’o: an advanced analysis and visualization platform for ‘omics data. PeerJ 2015;3:e1319.

38. Page AJ, Cummins CA, Hunt M, Wong VK, Reuter S, et al. Roary: rapid large-scale prokaryote pan genome analysis. Bioinformatics 2015;31:3691–3693.

39. Bortolaia V, Kaas RS, Ruppe E, Roberts MC, Schwarz S, et al. ResFinder 4.0 for predictions of phenotypes from genotypes. J Antimicrob Chemother 2020;75:3491–3500.

40. Seemann T. tseemann/abricate. https://github.com/tseemann/abricate (2026, accessed 21 April 2026).

41. Rafailidis PI, Kofteridis D. Proposed amendments regarding the definitions of multidrug-resistant and extensively drug-resistant bacteria. Expert Rev Anti Infect Ther 2022;20:139–146.

42. Robertson J, Nash JHE. MOB-suite: software tools for clustering, reconstruction and typing of plasmids from draft assemblies. Microb Genomics 2018;4:e000206.

43. Carattoli A, Hasman H. PlasmidFinder and In Silico pMLST: Identification and Typing of Plasmid Replicons in Whole-Genome Sequencing (WGS). In: de la Cruz F (editor). Horizontal Gene Transfer: Methods and Protocols. New York, NY: Springer US. pp. 285–294.

44. Redondo-Salvo S, Bartomeus-Peñalver R, Vielva L, Tagg KA, Webb HE, et al. COPLA, a taxonomic classifier of plasmids. BMC Bioinformatics 2021;22:390.

45. Xie Z, Tang H. ISEScan: automated identification of insertion sequence elements in prokaryotic genomes. Bioinformatics 2017;33:3340–3347.

46. Wishart DS, Han S, Saha S, Oler E, Peters H, et al. PHASTEST: faster than PHASTER, better than PHAST. Nucleic Acids Res 2023;51:W443–W450.

47. Pfeifer E, Rocha EPC. Phage-plasmids promote recombination and emergence of phages and plasmids. Nat Commun 2024;15:1545.

48. Li P, Lin H, Mi Z, Xing S, Tong Y, et al. Screening of Polyvalent Phage-Resistant Escherichia coli Strains Based on Phage Receptor Analysis. Front Microbiol;10. Epub ahead of print 18 April 2019. DOI: 10.3389/fmicb.2019.00850.

49. Liu L, Zhao D, Ye L, Zhan T, Xiong B, et al. A programmable CRISPR/Cas9-based phage defense system for Escherichia coli BL21(DE3). Microb Cell Factories 2020;19:136.

50. Chen M, Zhang L, Xin S, Yao H, Lu C, et al. Inducible Prophage Mutant of Escherichia coli Can Lyse New Host and the Key Sites of Receptor Recognition Identification. Front Microbiol;8. Epub ahead of print 1 February 2017. DOI: 10.3389/fmicb.2017.00147.

51. Néron B, Littner E, Haudiquet M, Perrin A, Cury J, et al. IntegronFinder 2.0: Identification and Analysis of Integrons across Bacteria, with a Focus on Antibiotic Resistance in Klebsiella. Microorganisms 2022;10:700.

52. Botelho J. Defense systems are pervasive across chromosomally integrated mobile genetic elements and are inversely correlated to virulence and antimicrobial resistance. Nucleic Acids Res 2023;51:4385–4397.

53. Néron B, Denise R, Coluzzi C, Touchon M, Rocha EPC, et al. MacSyFinder v2: Improved modelling and search engine to identify molecular systems in genomes. Peer Community J;3. Epub ahead of print 24 March 2023. DOI: 10.24072/pcjournal.250.

54. Tesson F, Hervé A, Mordret E, Touchon M, d’Humières C, et al. Systematic and quantitative view of the antiviral arsenal of prokaryotes. Nat Commun 2022;13:2561.

55. Doron S, Melamed S, Ofir G, Leavitt A, Lopatina A, et al. Systematic discovery of antiphage defense systems in the microbial pangenome. Science 2018;359:eaar4120.

56. Thiaville JJ, Kellner SM, Yuan Y, Hutinet G, Thiaville PC, et al. Novel genomic island modifies DNA with 7-deazaguanine derivatives. Proc Natl Acad Sci 2016;113:E1452–E1459.

57. Darracq B, Littner E, Brunie M, Bos J, Kaminski PA, et al. Sedentary chromosomal integrons as biobanks of bacterial antiphage defense systems. Science 2025;388:eads0768.

58. Getz LJ, Fairburn SR, Vivian Liu Y, Qian AL, Maxwell KL. Integrons are anti-phage defence libraries in Vibrio parahaemolyticus. Nat Microbiol 2025;10:724–733.

59. Horesh G, Fino C, Harms A, Dorman MJ, Parts L, et al. Type II and type IV toxin–antitoxin systems show different evolutionary patterns in the global Klebsiella pneumoniae population. Nucleic Acids Res 2020;48:4357–4370.

60. Rocha EPC, Bikard D. Microbial defenses against mobile genetic elements and viruses: Who defends whom from what? PLOS Biol 2022;20:e3001514.

61. Pinilla-Redondo R, Shehreen S, Marino ND, Fagerlund RD, Brown CM, et al. Discovery of multiple anti-CRISPRs highlights anti-defense gene clustering in mobile genetic elements. Nat Commun 2020;11:5652.

62. Tesson F, Huiting E, Wei L, Ren J, Johnson M, et al. Exploring the diversity of anti-defense systems across prokaryotes, phages and mobile genetic elements. Nucleic Acids Res 2025;53:gkae1171.

63. Gillings M, Boucher Y, Labbate M, Holmes A, Krishnan S, et al. The Evolution of Class 1 Integrons and the Rise of Antibiotic Resistance. J Bacteriol 2008;190:5095–5100.

64. Yao Y, Maddamsetti R, Weiss A, Ha Y, Wang T, et al. Intra- and interpopulation transposition of mobile genetic elements driven by antibiotic selection. Nat Ecol Evol 2022;6:555–564.

65. Samuel B, Mittelman K, Croitoru SY, Ben Haim M, Burstein D. Diverse anti-defence systems are encoded in the leading region of plasmids. Nature 2024;635:186–192.

66. Hernández-García M, Pérez-Viso B, Francisco CN-S, Baquero F, Morosini MI, et al. Intestinal co-colonization with different carbapenemase-producing Enterobacterales isolates is not a rare event in an OXA-48 endemic area. eClinicalMedicine 2019;15:72–79.

67. Maechler F, Schwab F, Hansen S, Fankhauser C, Harbarth S, et al. Contact isolation versus standard precautions to decrease acquisition of extended-spectrum β-lactamase-producing Enterobacterales in non-critical care wards: a cluster-randomised crossover trial. Lancet Infect Dis 2020;20:575–584.

68. Kolmogorov M, Yuan J, Lin Y, Pevzner PA. Assembly of long, error-prone reads using repeat graphs. Nat Biotechnol 2019;37:540–546.

69. Wick RR, Judd LM, Gorrie CL, Holt KE. Unicycler: Resolving bacterial genome assemblies from short and long sequencing reads. PLOS Comput Biol 2017;13:e1005595.

70. Wick RR, Schultz MB, Zobel J, Holt KE. Bandage: interactive visualization of de novo genome assemblies. Bioinformatics 2015;31:3350–3352.

71. Maiden MCJ, van Rensburg MJJ, Bray JE, Earle SG, Ford SA, et al. MLST revisited: the gene-by-gene approach to bacterial genomics. Nat Rev Microbiol 2013;11:728–736.

72. Seemann T. Prokka: rapid prokaryotic genome annotation. Bioinformatics 2014;30:2068–2069.

73. Katz LS, Griswold T, Morrison SS, Caravas JA, Zhang S, et al. Mashtree: a rapid comparison of whole genome sequence files. J Open Source Softw 2019;4:10.21105/joss.01762.

74. Letunic I, Bork P. Interactive Tree of Life (iTOL) v6: recent updates to the phylogenetic tree display and annotation tool. Nucleic Acids Res 2024;52:W78–W82.

75. Danecek P, Bonfield JK, Liddle J, Marshall J, Ohan V, et al. Twelve years of SAMtools and BCFtools. GigaScience 2021;10:giab008.

76. Li H. Aligning sequence reads, clone sequences and assembly contigs with BWA-MEM. Epub ahead of print 26 May 2013. DOI: 10.48550/arXiv.1303.3997.

77. Feldgarden M, Brover V, Gonzalez-Escalona N, Frye JG, Haendiges J, et al. AMRFinderPlus and the Reference Gene Catalog facilitate examination of the genomic links among antimicrobial resistance, stress response, and virulence. Sci Rep 2021;11:12728.

78. Sherry NL, Horan KA, Ballard SA, Gonҫalves da Silva A, Gorrie CL, et al. An ISO-certified genomics workflow for identification and surveillance of antimicrobial resistance. Nat Commun 2023;14:60.

79. Couvin D, Bernheim A, Toffano-Nioche C, Touchon M, Michalik J, et al. CRISPRCasFinder, an update of CRISRFinder, includes a portable version, enhanced performance and integrates search for Cas proteins. Nucleic Acids Res 2018;46:W246–W251.

